# Temporal characteristics of hemodynamic responses during active and passive hand movements in schizophrenia spectrum disorder

**DOI:** 10.1101/2025.05.15.653451

**Authors:** Harun A. Rashid, Tilo Kircher, Benjamin Straube

## Abstract

In healthy individuals active compared to passive movements exhibit earlier neural processing, reflected by more positive contrast estimates of the first-order temporal derivative (TD) of the hemodynamic response function (HRF) in functional MRI (fMRI) analyses. This temporal advantage may play a critical role in self-other distinction. However, whether Schizophrenia Spectrum Disorder (SSD) is associated with deficits in sensory-motor predictive mechanisms that influence this earlier processing remains unknown.

Patients with SSD (n = 20) and healthy control subjects (HC; n = 20) performed active and passive hand movements, while detected delays in video feedback of their own or another person’s hand movement. 3T fMRI data was recorded during the task. To assess response dynamics, we applied the TD to examine timing and the second-order dispersion derivative (DD) to evaluate duration of the HRF.

Compared to HC, patients with SSD exhibited delayed BOLD activation during active vs. passive movements in the right caudate nucleus, lobule VIII of right cerebellar hemisphere, left superior temporal gyrus, left postcentral gyrus, left thalamus, and left putamen/insula. For active movement with own hand feedback, HC showed earlier activation in the bilateral putamen and insula, whereas patients with SSD exhibited earlier activation in the left precentral gyrus, supplementary motor area, and postcentral gyrus.

Delayed BOLD responses in patients with SSD, particularly in the right cerebellar lobule VIII, bilateral putamen and insula, suggest impaired predictive mechanisms affecting feedback monitoring. These delays may contribute to disturbances in the sense of agency and self-action awareness, potentially underpinning core symptoms of SSD.

## Introduction

The core symptoms of schizophrenia spectrum disorder (SSD) are hallucinations and passivity symptoms. Patients with SSD often characterised by imprecise awareness about willed action, such as uncertain whether the willingness to initiate action planning and action initiation is their own. In this regard, impairments in patients with SSD have been demonstrated in sensory-motor feedback integration,^1–5^ action-consequences monitoring,^1,2,6–11^ self-other distinction in the fine-tune gripping and hand movement,^12–17^ feeling of agency control,^2,18–25^ and prediction of movement-related action consequence.^26–34^ While impairments in temporal dynamics of predictive processing might impair the brain’s ability to differentiate self-generated from externally-generated events^35^, evidence from fMRI studies in this domain is –to our knowledge-missing.

A functional MRI (fMRI) study in healthy subjects suggested that active hand movement shows earlier blood oxygenation level dependent (BOLD) response dynamics compared to passive hand movement.^36^ It has been suggested that efference copy (EC) related predictive mechanisms in active movement conditions might be responsible for this earlier activation.^36^ Similarly, an electroencephalography (EEG) study with speech articulation task results suggested that movement preparation related neural processes begin before stimulus onset.^37,38^ These neural processes develop corollary discharges (CD), which may include awareness of action-willingness and a general preparatory thought about cognitive-motor related neural preparation. CD subsequently forms motor commands specific to the sensorimotor task, and the copy of these commands from CD is known as EC.^37,38^ EC as an internally predicted representation of the tasks and their feedback, potentially facilitates brain activation patterns, which interacts with perception formation areas when feedback integration is processed.^36,37^ In this regard, fMRI studies reported reduced BOLD activation patterns in active compared to externally forced hand movement, and it is often suggested that these patterns are impaired in SSD patients.^2,39–44^ The awareness of control about task-intention, the timing of preparation initiation, CD, and EC could thus be the primary factors affecting the earlier or later temporal dynamics of BOLD responses. However, how active and passive hand movements induce earlier or later neural activation patterns in patients with SSD are yet to be explored.

Previously, we explored how neural activation patterns during hand movement preparation and execution differed and found that patients with SSD showed reduced neural activation during movement preparation, but not execution.^45^ For patients with SSD psychomotor slowing had been reported.^46,47^ The psychomotor slowing and impaired preparation, thus, might lead to slower or delayed neural processing. Therefore, we now applied an approach recently used in healthy subjects showing earlier neural processing of hand movement feedback for active compared to passive movements,^36^ to explore whether and how the timing of the activation patterns for hand movement-feedback processing is impaired in patients with SSD. This approach considers the temporal dynamics of BOLD responses by including the temporal derivative (TD) for time and the dispersion derivative (DD) for duration of the canonical hemodynamic response function (HRF) of BOLD responses in the analyses model. We expected that deficits in action-feedback monitoring might be reflected in the altered temporal dynamics of BOLD responses from self-generated (active) hand movement. Especially, we expect reduced differences between active and passive movements in patients with SSD due to a reduced active advantage (i.e. delayed processing in active conditions) in areas such as the supplementary motor area, dorsomedial prefrontal cortex, cerebellum, putamen and the insula. Especially the insula is not only involved in movement preparation,^45^ but also in the processing of multimodal sensory signals, and may involve in the external-internal subjective awareness.^48^

## Materials and Methods

### Participants

In this study, fMRI data from 20 HC and 20 patients with SSD (19 with schizophrenia and one with schizoaffective disorder [SAD]) were included. All subjects were right-handed. HC group (15 male; age: 38.4 ± 9.9 years old), patients with SSD (4 female; age: 38.5 ± 8.5 years). Patients with SSD were matched in terms of their age, sex, and attained highest educational degree. The detailed demographic and clinical characteristics, as well as additional information about the subjects, groups, stimuli, and equipment used in this study, have been reported in a previous publication.^28,49^ The patients were largely oligosymptomatic during the time of the experiment reflected in the Scale for the assessment of positive symptoms (SAPS).^50^ However, compared to HC patients with SSD showed a significantly higher SAPS score (12.1 ± 9.8): delusions (5.6 ± 5.1), delusions of reference (1.1 ± 1.2), delusions of being controlled (0.5 ± 0.9), residual positive symptoms (4.2 ± 5.1).^45^ This study was evaluated and approved by the local committee. Participants provided informed consent before the test measurement and were compensated for their participation. For further information see Table 1 from the previous study.^45^

**Table 1:**
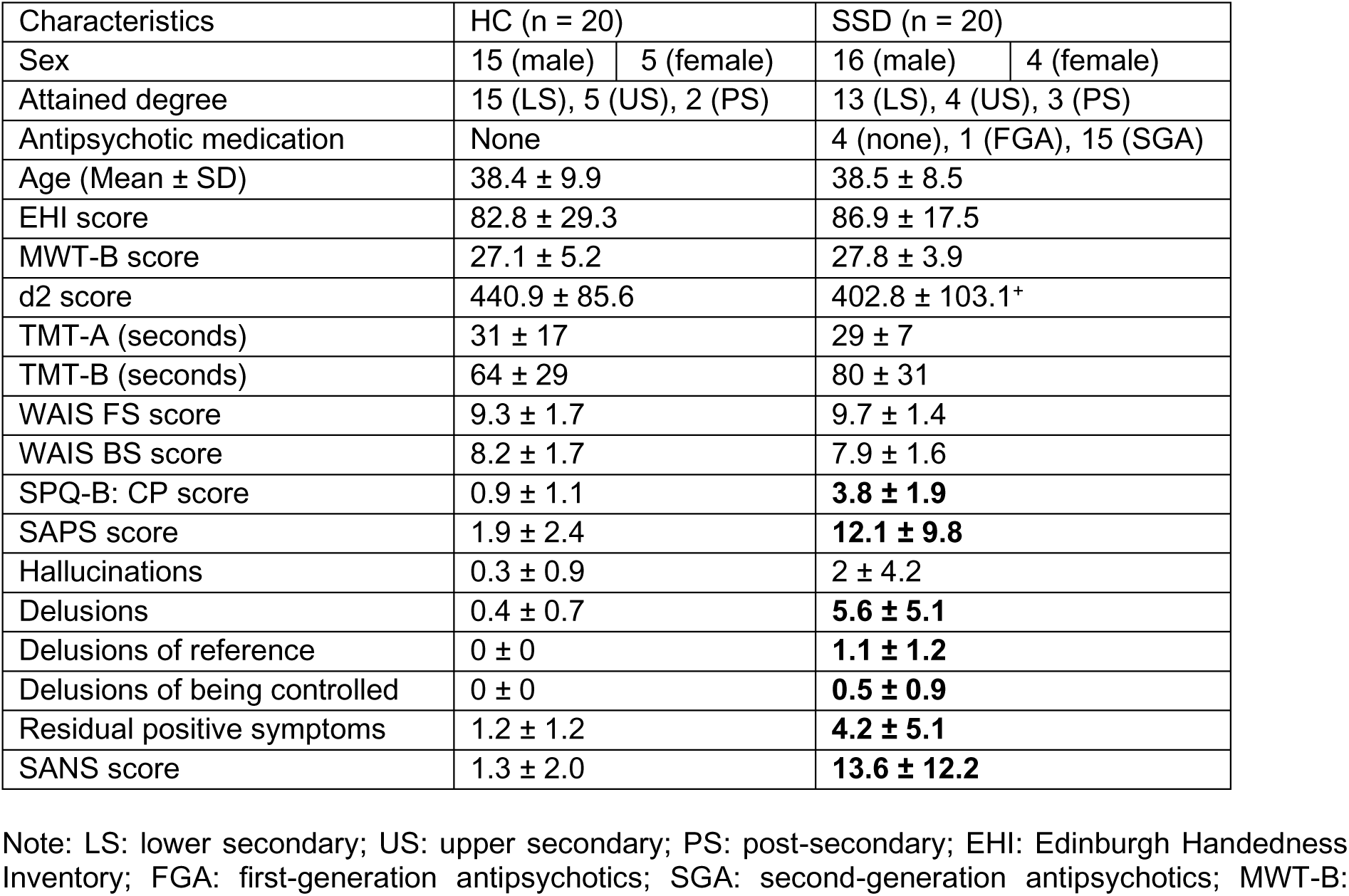
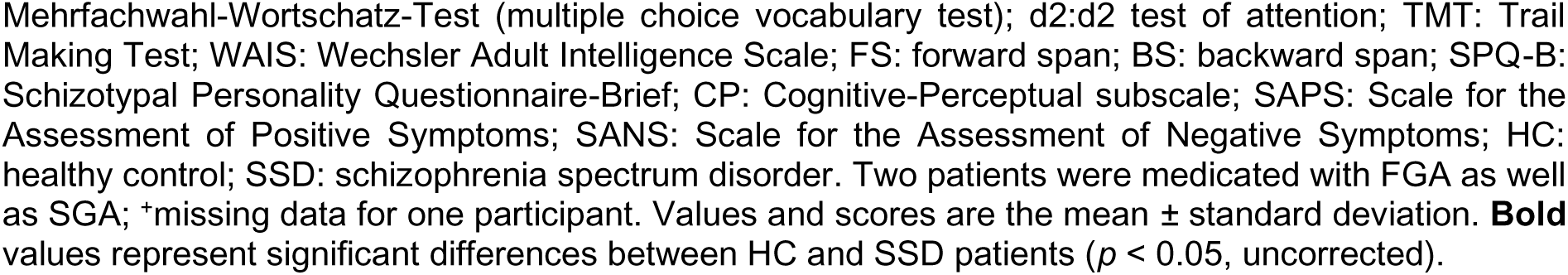
Demographic and clinical characteristics (from ^45^)

### Stimuli and Equipment

A custom-made MR-compatible passive movement device (PMD) was used, by gripping its handle hand could be moved from the left (home position) to the right end and back to the left home position through a circular arc (central angle: ∼30 degrees; trajectory: about 5.5 cm). Initiating planning and the execution of hand movement could be self-generated or moved by the PMD connected with an air pressure controller. Inside the MRI, the PMD was placed on the right thigh to facilitate right-hand movement. Handle position within ∼30° (movement trajectory) and direction of the hand movement were recorded via light emitting and detecting optical fibre cables integrated into the PMD.

The participants hands were simultaneously recorded via a high-speed camera (MRC High Speed, MRC Systems GmbH, Heidelberg, Germany; refresh rate: 4 ms) and displayed recorded video onto the screen (refresh rate: 60 Hz). During the experiment in the scanner, these visual feedback from the monitor was shown onto a tilted mirror. In the other 50% of trials, a similarly positioned pre-recorded hand from a person of the opposite sex were displayed, moving following their actual hand movement on the PMD. The video feedback was displayed with 6 different delays (0, 83, 167, 250, 334, 417 ms + internal setup delay of 43 ms) from the actual hand movement. These delays correspond to the screen’s refresh rate (0, 5, 10, 15, 20, and 25 frames at 60 Hz). Custom-written software on a computer (Intel® Core™ i5-4,570 CPU, 3.20 GHz, 4 GB RAM, AMD Radeon HD8570D Graphics Card, 32-bit operating system, Windows 7 Professional [Microsoft Corporation, 2009]) was used to control the setup.

### Experimental Design

A mixed factorial design with the within-subject factors, movement execution (active and passive), video feedback (own and other), as well as a between-subjects factor group (patients with SSD and HC) was used. This results in four movements and feedback conditions: self-active, self-passive, other-active, and other-passive.

There were 2 fMRI sessions with two runs of 48 trials in each session. Each run began with cues ("Active" or "Passive"), instructing active or passive blocks having sequential 24 trials in each block. Each trial initiates with a "Ready" cue, followed by a randomized video display of the hand ("self" or "other") grasping the knob/handle of PMD. In active trials, participants could prepare and move their hand actively during the displayed video. While in passive trials, hand movement was programmed to be automatically initiated by the PMD with 500 ms (additionally, internal delay of compressor and PMD) delay after the camera onset; to keep the similarity in the movement one timing between the active and passive trials. . Here equally and randomly distributed 6 different delays between 0 - 417 ms were applied between video feedback and actual hand movement. Subsequently, a "Delay?" cue appeared to respond to perceived delay in the feedback via button press; left hand’s index (no) or middle finger to say yes. Trials concluded with a black screen of inter-trial interval, randomized between 2000-5000 ms.

### Procedure

Participants were familiarized with the task and setup in a preparatory behavioural session. Where participants were sitting upright in front of a computer screen. The preparatory session was followed by fMRI sessions with 2 runs (each having 48 trials.). Inside the MRI, subjects were lying in a supine position, while the PMD was placed onto the right thigh. Participants were directed to keep gripping the PMD handle with their right hand using the index finger and thumb for the upper part, and the rest fingers for the lower part of the handle. Hand movements consisted of an extension from the left to the right and move it back from the right to the left home position. In the active condition, the subjects were asked, and trained in the preparatory session to complete each movement in about 1500 ms in order to make it consistent with the passive condition, where the subjects just held the handle and the PMD executed the hand movement. At the end of the experiment, all subjects filled out a post experiment questionnaire.

### Functional Data Acquisition

Data was acquired using a 3 T Magnetom Trio Tim scanner (Siemens, Erlangen, Germany) with a 12-channel head coil. For functional data, a T2*-weighted gradient-echo echoplanar imaging sequence (TR: 1,650 ms, TE: 25 ms, flip angle: 70°) was procured. In each run 330 volumes were collected in a descending order, each covering 34 axial brain slices (matrix: 64 × 64, field of view [FoV]: 192 mm × 192 mm, slice thickness: 4 mm, voxel size: 3 mm × 3 mm × 4.6 mm [with a 0.6 mm gap]). Anatomical images were acquired using a T1-weighted MPRAGE sequence (TR: 1,900 ms, TE: 2.26 ms, flip angle: 9°) with a matrix of 256 × 256, FoV of 256 mm × 256 mm, slice thickness of 1 mm, and voxel size of 1 mm × 1 mm × 1.5 mm [including a 0.5 mm gap].

### Statistical Analysis

The standard procedures from Statistical Parametric Mapping (SPM12 version 7, Wellcome Trust Centre for Neuroimaging, University College London, UK) were implemented in MATLAB 2017a (The Mathworks Inc.) to perform statistical analyses of fMRI data. Correlation related analyses were performed using JASP (Jeffreys’s Amazing Statistics Program; version 0.18.3; JASP team, 2024). The preprocessing steps involved: Realignment, Coregistration between anatomical and functional images, Segmentation, Normalization to the Montreal Neurological Institute (MNI) standard space (resampled to a voxel size of 2 mm × 2 mm × 2 mm), and smoothing (8 mm × 8 mm × 8 mm full-width at half-maximum Gaussian kernel). The framewise displacement computed between consecutive volumes, only < 10% values exceeded 1 mm in any run. Here, the focus is on the fMRI contrast relevant to the complete movement video feedback period (4000 ms; see Figure 1).

**Figure 1:**
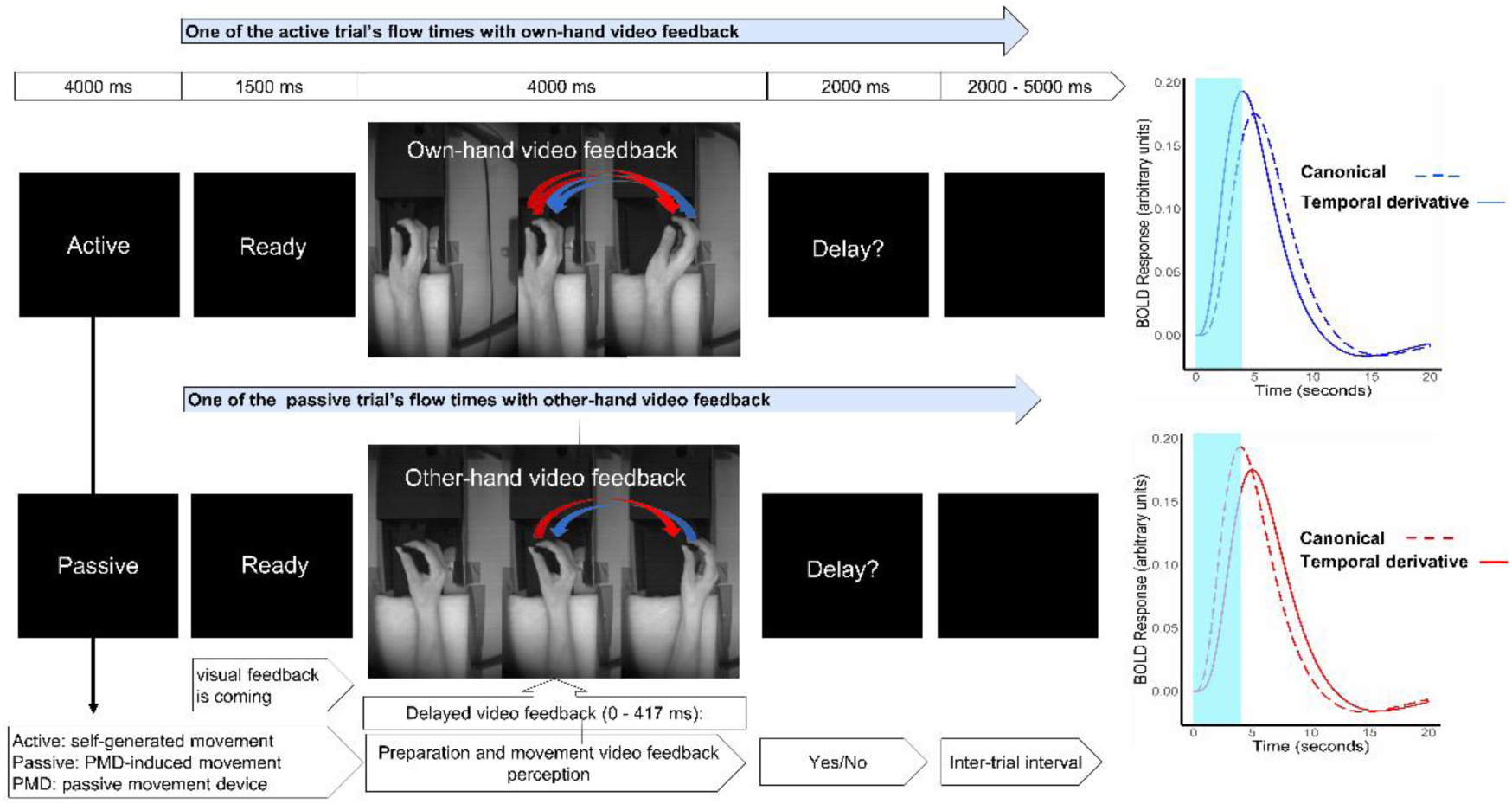
Experimental design. In the beginning of each run (48 trials), participants were instructed to perform hand movement by themselves when cued “Active” (24 trials), or to relax the hand but keep holding the PMD handle when cued “Passive” and let their hand be moved by the PMD. A trial commenced with a “Ready” cue, followed by visual feedback for movement planning and execution, culminating with a question “Delay?”. The black screen with variable duration (2000 - 5000 ms) is shown during inter trial interval. In the planning and preparation period, the hand video feedback was mostly static (1st hand) since the subject was instructed to move only to execute. The movement onset from the left (2^nd^ hand) to the right (3^rd^ hand), then the hand moves it back to the left (from 3rd hand to the 2^nd^ hand) position. In the video feedback, only the right-hand movement was displayed, here 3 different hand positions are shown just to visualize the whole process of planning and execution direction, separately. Considering male participants for this figure, the upper row shows the sequence of a trial with active and own-hand video feedback, while the lower row shows another trial with the passive and other hand image from the female. In the case of a female subject, the other hand video is shown from a male hand image. Self-other hand was displayed randomly across the trial and run. A video demonstration is available at http://doi.org/10.5281/zenodo.2621302. On the right, the expected active and passive movement specific canonical hemodynamic response function and temporal derivative of BOLD responses are shown (the 4000 ms of movement with feedback stimulus marked with light blue.

In these analyses, the acquired blood oxygenation level dependent (BOLD) response for each trial was modelled from the time of camera onset to camera offset. Head movement was further included as control regressor. Therefore, for each participant, 4 regressors were modelled as conditions of interest, cue, and question periods as well as the 6-motion parameter were included as a nuisance regressor into the General Linear Model (GLM). The regressors were convolved with the canonical hemodynamic response function (HRF). BOLD responses were further modelled by utilizing the first and second-order Taylor expansion of the canonical hemodynamic response function (HRF); thus, the first-order derivative concerning time (TD) and second-order derivative for the duration (DD) were included. TD reveals a distinction in the BOLD response timing, where DD reflects the duration of the BOLD response.^51^ This is then compared with canonical HRF, to perform statistical analyses to reveal the relative timing reflected in TD and duration in DD compared to canonical timing and duration.^51,52^ While the inclusion of TD improves sensitivity to the timing and onset dynamics of neural processes,^51,53^ it alone is insufficient to capture alterations in the shape or duration of the hemodynamic response. Incorporating DD enables the detection of differences in the duration of neural processes, particularly highlighting additional alterations identified in patients with SSD compared to HC. In our analyses, canonical HRF analyses remain the most sensitive for detecting variations in BOLD amplitude, while the addition of TD enhances detection of onset-related temporal dynamics. Similarly, adding DD improves sensitivity to detect durations of sustained neural activity but might attenuate the detection sensitivity,^54^ as we observed both on amplitude and timing variations. Accordingly, we first analysed HRF+TD to optimally detect timing-related group differences. To identify duration-related alterations, we conducted a second analysis using HRF+TD+DD.

To remove low-frequency noise from the time series a 128 s high pass filter was applied. Finally, we contrasted regressors of interest against the implicit baseline for each participant, and the resulting contrast estimates were entered into a group-level full factorial model. T and F contrast for the TD and DD were created in each participants data, for every regressor of movement feedback condition (self-active, self-passive, other-active, and other passive) in the first level. This is then used in the second-level analyses, yielding 3 contrasts per subject.

Here the identical cluster threshold of 104 voxels and cluster defining threshold of p < 0.005 uncorrected (Note: Family-Wise-Error (P_FWC-corrected)_ values are additionally provided in the Table 1, 2, and 3), as previously determined by Monte Carlo simulations,^28^ was applied to correct for multiple comparisons at P < 0.05. This ensures consistency with the previous analyses and results. Additionally, p-values for FWE cluster correction will be provided in Tables 1, 2, and 3. Coordinates are listed using standard MNI152 space. To explore activation timing and duration, we analysed the following T-contrasts within HC and SSD groups and compared them in conjunction analyses, between-group, and interaction contrast: active>passive; contrast for the commonalities: HC (active>passive) Ո SSD (active>passive); contrast for the group differences: HC (active>passive) > SSD (active>passive); interaction with group differences: [HC ((Selfact-Selfpas)-(Otheract-Otherpas))]>[SSD((Selfact-Selfpas)- (Otheract-Otherpas))]. The same analyses were conducted for passive>active contrast. For the condition and group-specific cluster and effect, contrast masking was implemented. To investigate the group-specific effect, HC (active>passive) > SSD (active>passive) masked by HC (active>passive); SSD (active>passive) > HC (active>passive) masked by SSD (active>passive); HC (passive>active) > SSD (passive>active) masked by HC (passive>active); for condition-specific effect, HC((Selfact-Selfpas)-(Otheract-Otherpas))>SSD((Selfact-Selfpas)-(Otheract-Otherpas)) masked by HC (Self-active) were analysed.

**Table 1:**
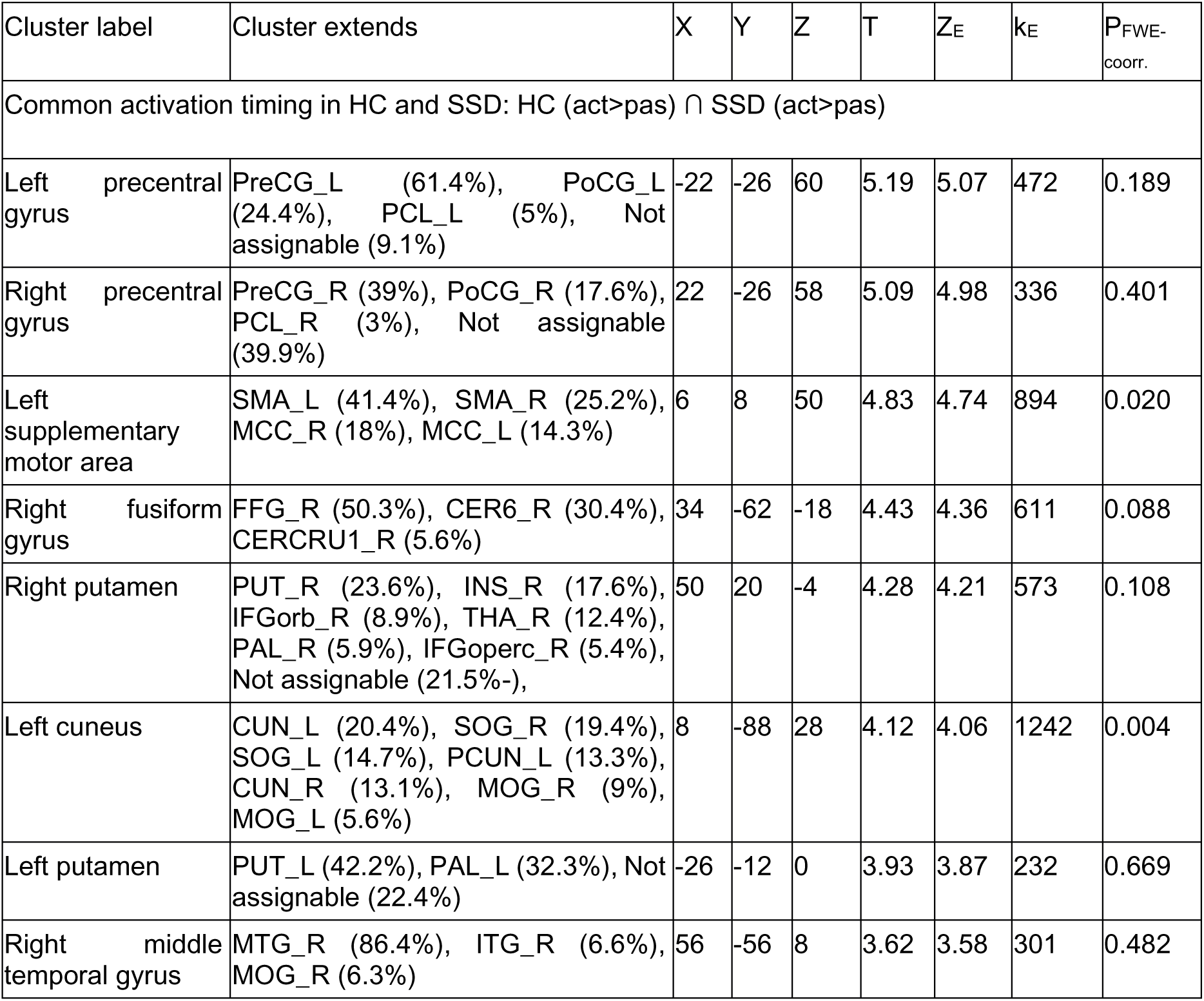
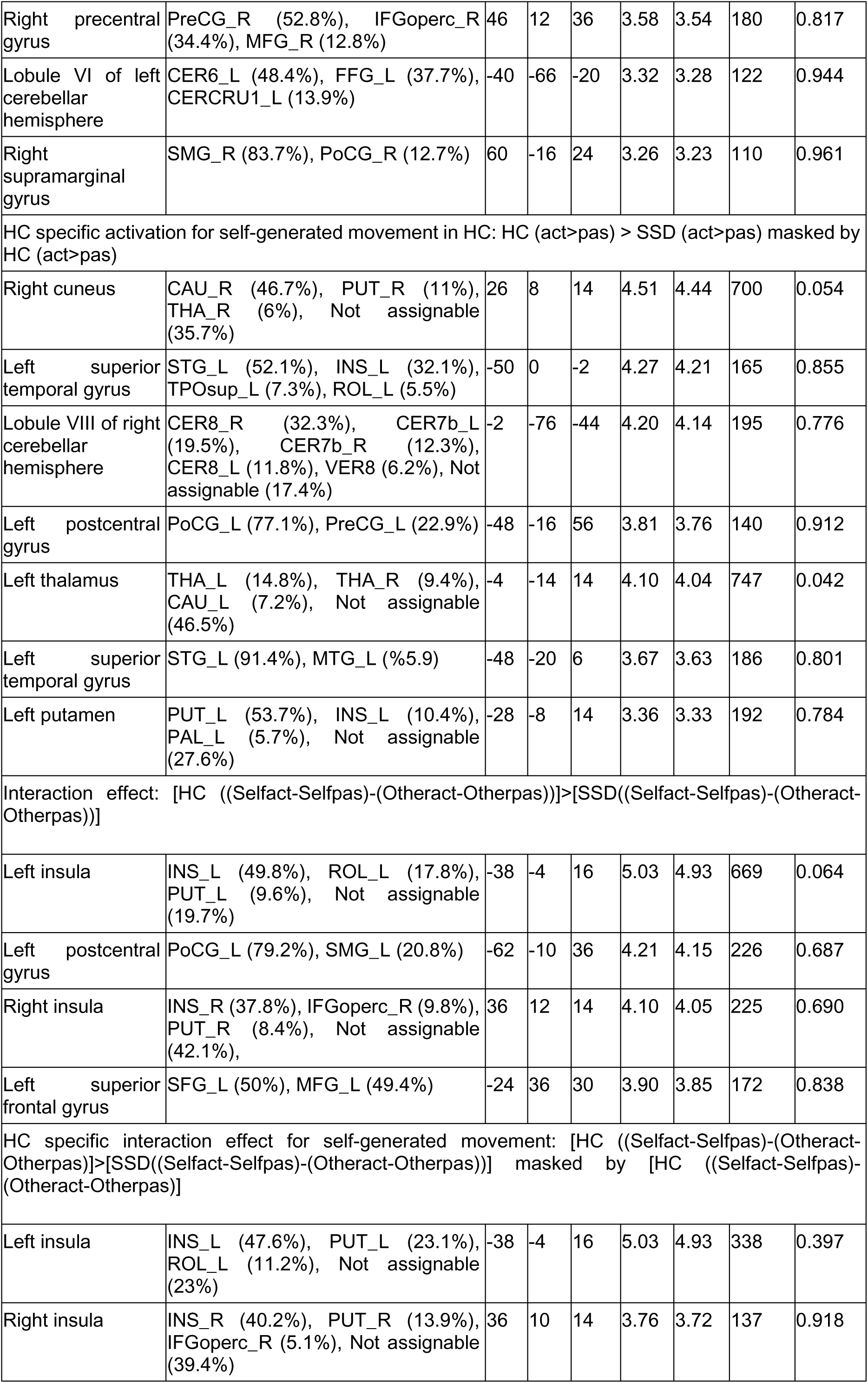
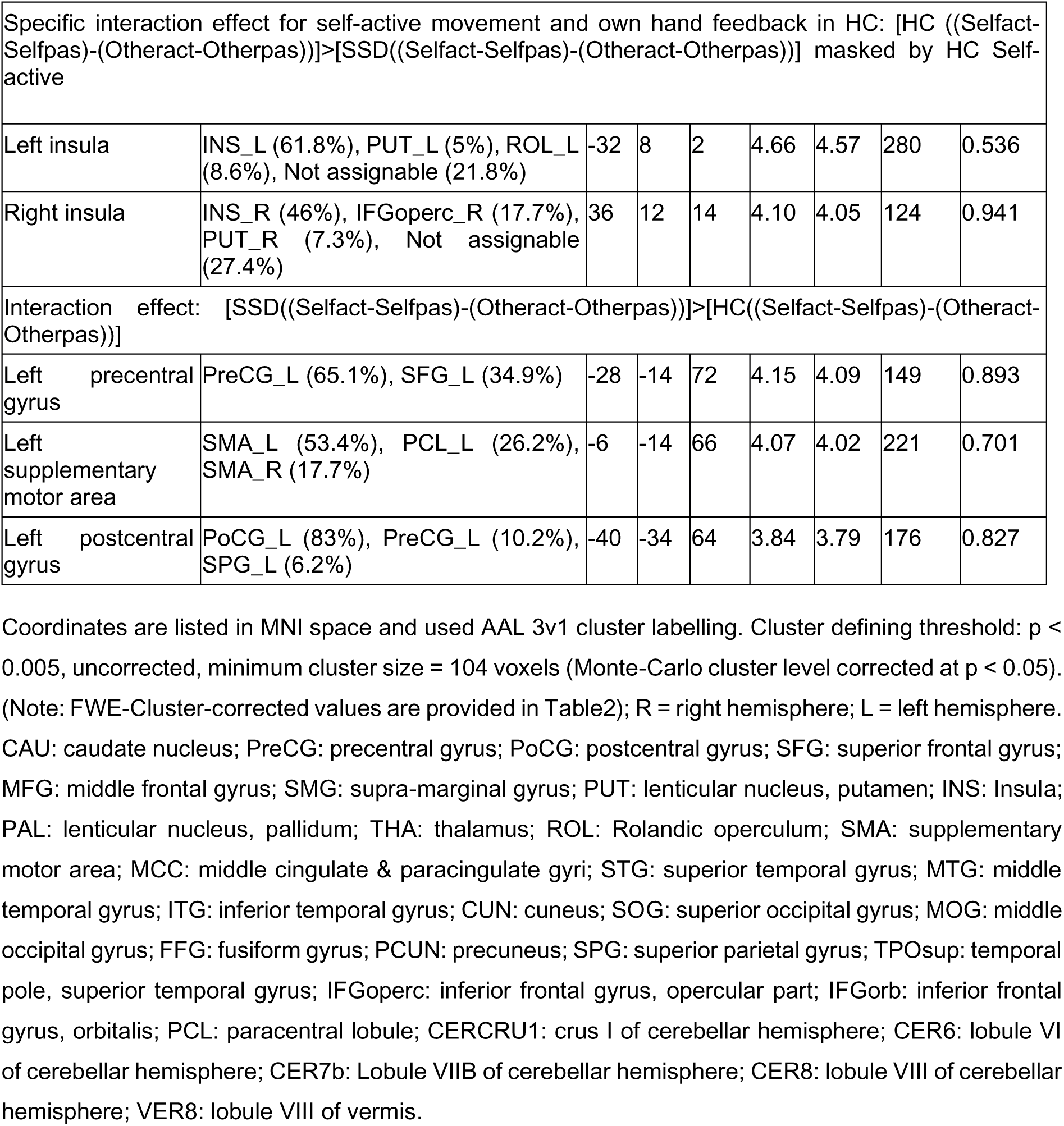
Results regarding activation timing from TD: active > passive.

| Cluster label | Cluster extends | X | Y | Z | T | Z<sub>E</sub> | k<sub>E</sub> | P<sub>FWE-coorr.</sub> |
| --- | --- | --- | --- | --- | --- | --- | --- | --- |
| Common activation timing in HC and SSD: HC (act>pas) ∩ SSD (act>pas) | | | | | | | | |
| Left precentral gyrus | PreCG_L (61.4%), PoCG_L (24.4%), PCL_L (5%), Not assignable (9.1%) | -22 | -26 | 60 | 5.19 | 5.07 | 472 | 0.189 |
| Right precentral gyrus | PreCG_R (39%), PoCG_R (17.6%), PCL_R (3%), Not assignable (39.9%) | 22 | -26 | 58 | 5.09 | 4.98 | 336 | 0.401 |
| Left supplementary motor area | SMA_L (41.4%), SMA_R (25.2%), MCC_R (18%), MCC_L (14.3%) | 6 | 8 | 50 | 4.83 | 4.74 | 894 | 0.020 |
| Right fusiform gyrus | FFG_R (50.3%), CER6_R (30.4%), CERCRU1_R (5.6%) | 34 | -62 | -18 | 4.43 | 4.36 | 611 | 0.088 |
| Right putamen | PUT_R (23.6%), INS_R (17.6%), IFGorb_R (8.9%), THA_R (12.4%), PAL_R (5.9%), IFGoperc_R (5.4%), Not assignable (21.5%-), | 50 | 20 | -4 | 4.28 | 4.21 | 573 | 0.108 |
| Left cuneus | CUN_L (20.4%), SOG_R (19.4%), SOG_L (14.7%), PCUN_L (13.3%), CUN_R (13.1%), MOG_R (9%), MOG_L (5.6%) | 8 | -88 | 28 | 4.12 | 4.06 | 1242 | 0.004 |
| Left putamen | PUT_L (42.2%), PAL_L (32.3%), Not assignable (22.4%) | -26 | -12 | 0 | 3.93 | 3.87 | 232 | 0.669 |
| Right middle temporal gyrus | MTG_R (86.4%), ITG_R (6.6%), MOG_R (6.3%) | 56 | -56 | 8 | 3.62 | 3.58 | 301 | 0.482 |
| Right precentral gyrus | PreCG_R (52.8%), IFGoperc_R (34.4%), MFG_R (12.8%) | 46 | 12 | 36 | 3.58 | 3.54 | 180 | 0.817 |
| Lobule VI of left cerebellar hemisphere | CER6_L (48.4%), FFG_L (37.7%), CERCRU1_L (13.9%) | -40 | -66 | -20 | 3.32 | 3.28 | 122 | 0.944 |
| Right supramarginal gyrus | SMG_R (83.7%), PoCG_R (12.7%) | 60 | -16 | 24 | 3.26 | 3.23 | 110 | 0.961 |
| HC specific activation for self-generated movement in HC: HC (act>pas) > SSD (act>pas) masked by HC (act>pas) |  |  |  |  |  |  |  |  |
| Right cuneus | CAU_R (46.7%), PUT_R (11%), THA_R (6%), Not assignable (35.7%) | 26 | 8 | 14 | 4.51 | 4.44 | 700 | 0.054 |
| Left superior temporal gyrus | STG_L (52.1%), INS_L (32.1%), TPOsup_L (7.3%), ROL_L (5.5%) | -50 | 0 | -2 | 4.27 | 4.21 | 165 | 0.855 |
| Lobule VIII of right cerebellar hemisphere | CER8_R (32.3%), CER7b_L (19.5%), CER7b_R (12.3%), CER8_L (11.8%), VER8 (6.2%), Not assignable (17.4%) | -2 | -76 | -44 | 4.20 | 4.14 | 195 | 0.776 |
| Left postcentral gyrus | PoCG_L (77.1%), PreCG_L (22.9%) | -48 | -16 | 56 | 3.81 | 3.76 | 140 | 0.912 |
| Left thalamus | THA_L (14.8%), THA_R (9.4%), CAU_L (7.2%), Not assignable (46.5%) | -4 | -14 | 14 | 4.10 | 4.04 | 747 | 0.042 |
| Left superior temporal gyrus | STG_L (91.4%), MTG_L (5.9%) | -48 | -20 | 6 | 3.67 | 3.63 | 186 | 0.801 |
| Left putamen | PUT_L (53.7%), INS_L (10.4%), PAL_L (5.7%), Not assignable (27.6%) | -28 | -8 | 14 | 3.36 | 3.33 | 192 | 0.784 |
| Interaction effect: [HC ((Selfact-Selfpas)-(Otheract-Otherpas))]>[SSD((Selfact-Selfpas)-(Otheract-Otherpas))] |  |  |  |  |  |  |  |  |
| Left insula | INS_L (49.8%), ROL_L (17.8%), PUT_L (9.6%), Not assignable (19.7%) | -38 | -4 | 16 | 5.03 | 4.93 | 669 | 0.064 |
| Left postcentral gyrus | PoCG_L (79.2%), SMG_L (20.8%) | -62 | -10 | 36 | 4.21 | 4.15 | 226 | 0.687 |
| Right insula | INS_R (37.8%), IFGoperc_R (9.8%), PUT_R (8.4%), Not assignable (42.1%), | 36 | 12 | 14 | 4.10 | 4.05 | 225 | 0.690 |
| Left superior frontal gyrus | SFG_L (50%), MFG_L (49.4%) | -24 | 36 | 30 | 3.90 | 3.85 | 172 | 0.838 |
| HC specific interaction effect for self-generated movement: [HC ((Selfact-Selfpas)-(Otheract-Otherpas))]>[SSD((Selfact-Selfpas)-(Otheract-Otherpas))] masked by [HC ((Selfact-Selfpas)-(Otheract-Otherpas))] |  |  |  |  |  |  |  |  |
| Left insula | INS_L (47.6%), PUT_L (23.1%), ROL_L (11.2%), Not assignable (23%) | -38 | -4 | 16 | 5.03 | 4.93 | 338 | 0.397 |
| Right insula | INS_R (40.2%), PUT_R (13.9%), IFGoperc_R (5.1%), Not assignable (39.4%) | 36 | 10 | 14 | 3.76 | 3.72 | 137 | 0.918 |
| Specific interaction effect for self-active movement and own hand feedback in HC: [HC ((Selfact-Selfpas)-(Otheract-Otherpas))]>[SSD((Selfact-Selfpas)-(Otheract-Otherpas))] masked by HC Self-active |  |  |  |  |  |  |  |  |
| Left insula | INS_L (61.8%), PUT_L (5%), ROL_L (8.6%), Not assignable (21.8%) | -32 | 8 | 2 | 4.66 | 4.57 | 280 | 0.536 |
| Right insula | INS_R (46%), IFGoperc_R (17.7%), PUT_R (7.3%), Not assignable (27.4%) | 36 | 12 | 14 | 4.10 | 4.05 | 124 | 0.941 |
| Interaction effect: [SSD((Selfact-Selfpas)-(Otheract-Otherpas))]>[HC((Selfact-Selfpas)-(Otheract-Otherpas))] |  |  |  |  |  |  |  |  |
| Left precentral gyrus | PreCG_L (65.1%), SFG_L (34.9%) | -28 | -14 | 72 | 4.15 | 4.09 | 149 | 0.893 |
| Left supplementary motor area | SMA_L (53.4%), PCL_L (26.2%), SMA_R (17.7%) | -6 | -14 | 66 | 4.07 | 4.02 | 221 | 0.701 |
| Left postcentral gyrus | PoCG_L (83%), PreCG_L (10.2%), SPG_L (6.2%) | -40 | -34 | 64 | 3.84 | 3.79 | 176 | 0.827 |

**Table 2:** Results regarding activation timing from TD: passive > passive.

| Cluster label | Cluster extend | X | Y | Z | T | Z<sub>E</sub> | k<sub>E</sub> | P<sub>FWE-coorr.</sub> |
| --- | --- | --- | --- | --- | --- | --- | --- | --- |
| Interaction effect: [HC((Selfpas-Selfact)-(Otherpas-Otheract))] > [SSD((Selfpas-Selfact)-(Otherpas-Otheract))] |  |  |  |  |  |  |  |  |
| Left paracentral lobule | PCL_L (12.4%), SMA_R (11.5%), PoCG_L (10.6%), PCUN_L (10.3%), SMA_L (8.4%), SFG_R (8.1%), SPG_R (6.9%), PreCG_R (6.9%), SFG_L (6.7%) | 6 | -2 | 76 | 4.71 | 4.62 | 3134 | < 0.001 |
| Left middle temporal gyrus | MTG_L (50.2%), Not assignable (48.5%) | -48 | -32 | -6 | 4.68 | 4.60 | 229 | 0.678 |
| Lobule VIII of left cerebellar hemisphere | CER8_L (59.4%), CER9_L (12.7%), Not assignable (24.2%) | -8 | -62 | -52 | 4.25 | 4.19 | 244 | 0.635 |
| Right parahippocampal gyrus | PHG_R (19.3%), HIP_R (19.3%), FFG_R (13.3%) | 32 | -46 | 0 | 3.93 | 3.88 | 181 | 0.814 |

Coordinates are listed in MNI space and used AAL 3v1 cluster labelling. Cluster defining threshold: $p < 0.005$ , uncorrected, minimum cluster size = 104 voxels (Monte-Carlo cluster level corrected at $p < 0.05$ ). (Note: FWE-Cluster-corrected values are provided in Table 2); R = right hemisphere; L = left hemisphere. PCL: paracentral lobule; SMA: supplementary motor area; PreCG: precentral gyrus; PoCG: postcentral gyrus; PCUN: precuneus; SFG: superior frontal gyrus; SPG: superior parietal gyrus; MTG: middle temporal gyrus; CER8: lobule VIII of cerebellar hemisphere; CER9: lobule IX of cerebellar hemisphere; HIP: hippocampus; PHG: parahippocampal gyrus; FFG: fusiform gyrus.

The name of the brain area is defined as the cluster name with the highest percent contribution to a cluster( using AAL 3v1 cluster labelling).^55^ The cluster extension of brain regions (>5% contribution) with their percentage of contribution are listed in Tables 1, 2, and 3. Only clusters were listed from which more than 50% percent voxel could be labelled. A cluster label with an MNI152 space coordinate (X Y Z) is used to interpret the result. Thus, all of the results are shown here in the MNI152 space’s coordinate. Latency functions (LF) relative to the canonical hemodynamic response function (HRF) of BOLD responses were assessed using methods described in the previous studies ^36,52^

### Exploratory Correlation Analyses

Here, we investigated how BOLD response timings and durations reflected by eigenvariates of the clusters in SSD patients are associated with negative and positive symptoms. The positive symptoms were scored by subscales of hallucinations (items 1–7), delusions of reference (item 14), delusions of being controlled (item 15)), and residual positive symptoms (items 21–35) of the assessment of positive symptoms (SAPS).^50^ The negative symptoms were assessed using related subscales of affective flattening or blunting (items 1–8), alogia (items 9–13), avolition/apathy (items 14–17), anhedonia/asociality (items 18–22), and attention (items 23–25 of the scale for the assessment of negative symptoms (SANS).^56^ The correlation of positive and negative symptom scores with neural activations timings (extracted eigenvariates) in active movement (bilateral insula/putamen) with own hand video feedback were performed. To explore specificity regarding the positive symptoms, partial correlations were performed by partialling out total SANS score. Analyses were performed using JASP (Jeffreys’s Amazing Statistics Program; version 0.18.3; JASP team, 2024).

### Data availability

The information about data supporting this study’s findings will be published upon the acceptance

## Results

### fMRI results

#### Group commonalities in the temporal dynamics of BOLD responses (active>passive)

Patients with SSD, compared to HC, exhibited largely comparable temporal dynamics of BOLD responses in clusters primarily comprising the bilateral precentral gyrus, bilateral putamen, right fusiform gyrus, right middle temporal gyrus, right supramarginal gyrus, left supplementary motor area, left cuneus, and lobule VI of left cerebellar hemisphere, as shown in (Table 1, Figure 2c).

**Figure 2:**
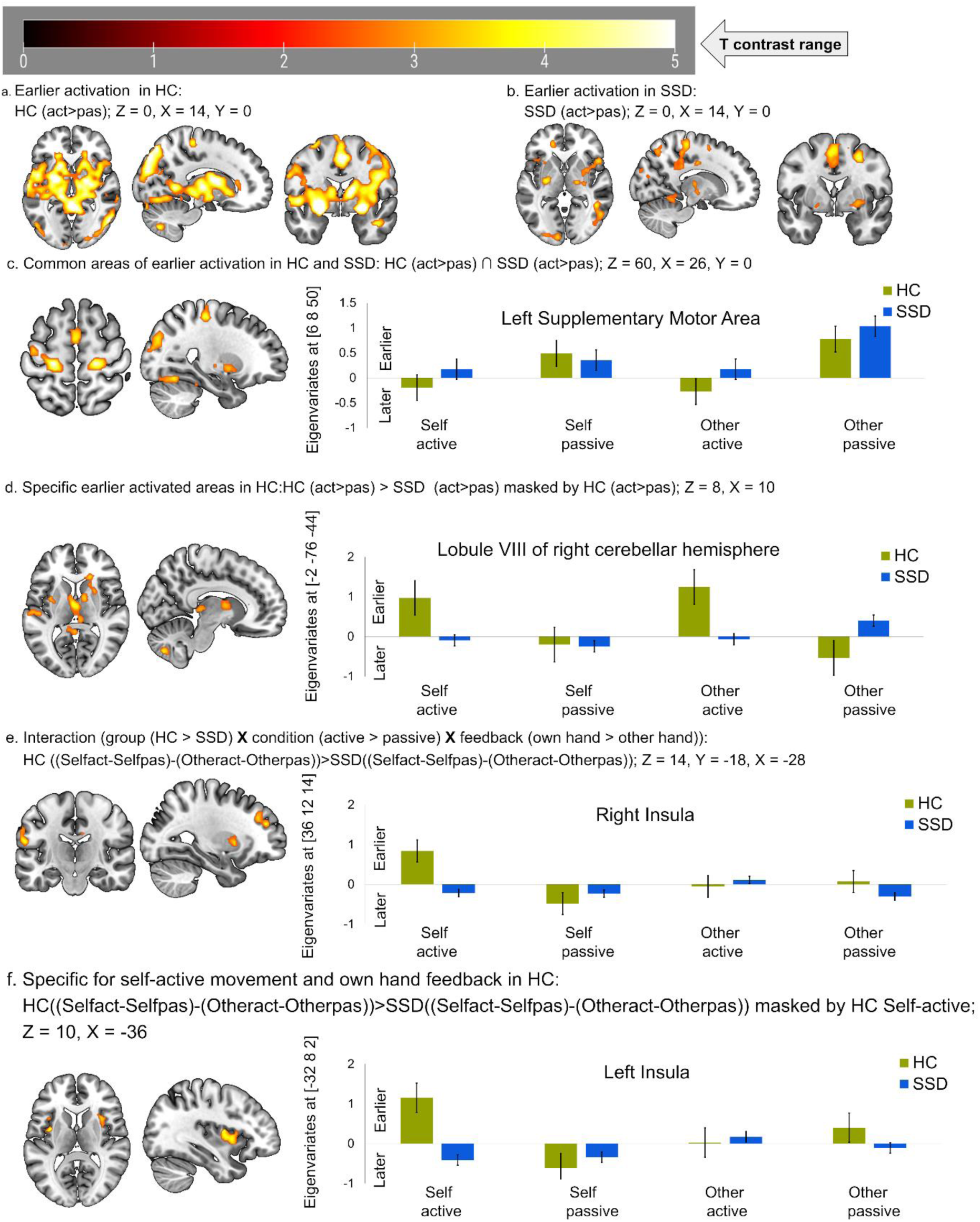
Timing of neural activation in active compared to passive condition. (a) earlier activation in healthy control (HC) shown at Z = 0, Y = 0, X = 14; (b) earlier activation in schizophrenia spectrum disorder (SSD) shown at Z = 0, Y = 0, X = 14; (c). common earlier activated brain areas between HCand SSD at Z = 60, Y = 0, X = 26, (d) earlier activation specific to HC compared to SSD shown at Z = 8, X = 10; (e) interaction related earlier activation in HC compared SSD patients shown at Z = 14, Y = −18, X = −28; f. active with own hand feedback specific earlier activated brain area in HC compared to SSD patients shown at Z = 10, X = −36. Positive eigenvariates reflect earlier activation while negative reflects later activation timing in the bar graph. HC: healthy control, (n= 20); SSD: schizophrenia spectrum disorder, (n = 20).

#### Group differences in the temporal dynamics of BOLD responses (active>passive)

We analysed whether patients with SSD differed in the active (compared to passive) movement specific earlier activation patterns observed in HC by applying inclusive masking to the group interaction with the HC contrast. Compared to HC, SSD patients exhibited delayed BOLD activation patterns in several key clusters mainly consisting the right caudate nucleus, lobule VIII of the right cerebellar hemisphere, left superior temporal gyrus, left postcentral gyrus, left thalamus, and left putamen during active compared to passive hand movements (Table 1, Figure 2d)

#### Interaction analysis

A three-factor interaction of group (HC>SSD), movement condition (active > passive), and video feedback (own hand > other hand) revealed clusters comprising mainly bilateral insula, left postcentral gyrus, and left superior frontal gyrus (Table 1, Figure 2e). In contrast to HC, SSD patients demonstrated relatively earlier activation in the left precentral gyrus, left supplementary motor area, and left postcentral gyrus during active relative to passive movements with their own hand feedback (Table 1).

#### Group timing differences during active movements with own-hand feedback

We further investigated whether specific (sub-) cortical regions exhibited timing differences during active movement with own hand feedback. Interestingly, HC showed earlier activation than SSD patients for active vs passive movements with own hand feedback, within both hemispheres the clusters primarily comprising the bilateral putamen and bilateral insula (Table 1, Figure 2f).

#### Group differences in temporal dynamics of BOLD responses (passive > active)

Although contrast (passive > active) revealed distinct clusters within the group of HC and SSD patients, no commonalities were found with similarly activated timing of BOLD responses. Also, no group differences were found. Thus, group differences in earlier activation patterns are specific to active compared to passive movement.

#### Duration of BOLD activation (active > passive)

The conjunction analyses revealed no overlap in activation for the active > passive contrast regarding durations (DD) between HC and SSD patients. Main interaction of group (HC > SSD), movement condition (active > passive), and feedback (own hand > other hand) analyses revealed clusters comprising primarily the Left precentral gyrus, right caudate nucleus, left superior parietal gyrus, right middle temporal gyrus, right precuneus, right parahippocampal gyrus, and left middle frontal gyrus (Table 3, Figure 3a). Additionally, to explore these differences in areas showing a specific pattern in HC, the interaction of group (HC > SSD), movement condition (active > passive), and feedback (own hand > other hand) were masked by either the interaction in HC [movement condition (active > passive) and feedback (own hand > other hand)] or masked by active movement with own hand feedback in HC. Both of these analyses revealed clusters mainly consisting Left precentral gyrus, right caudate nucleus, left superior parietal gyrus, and lobule VI of left cerebellar hemisphere (Table 3, Figure 3b).

**Table 3:**
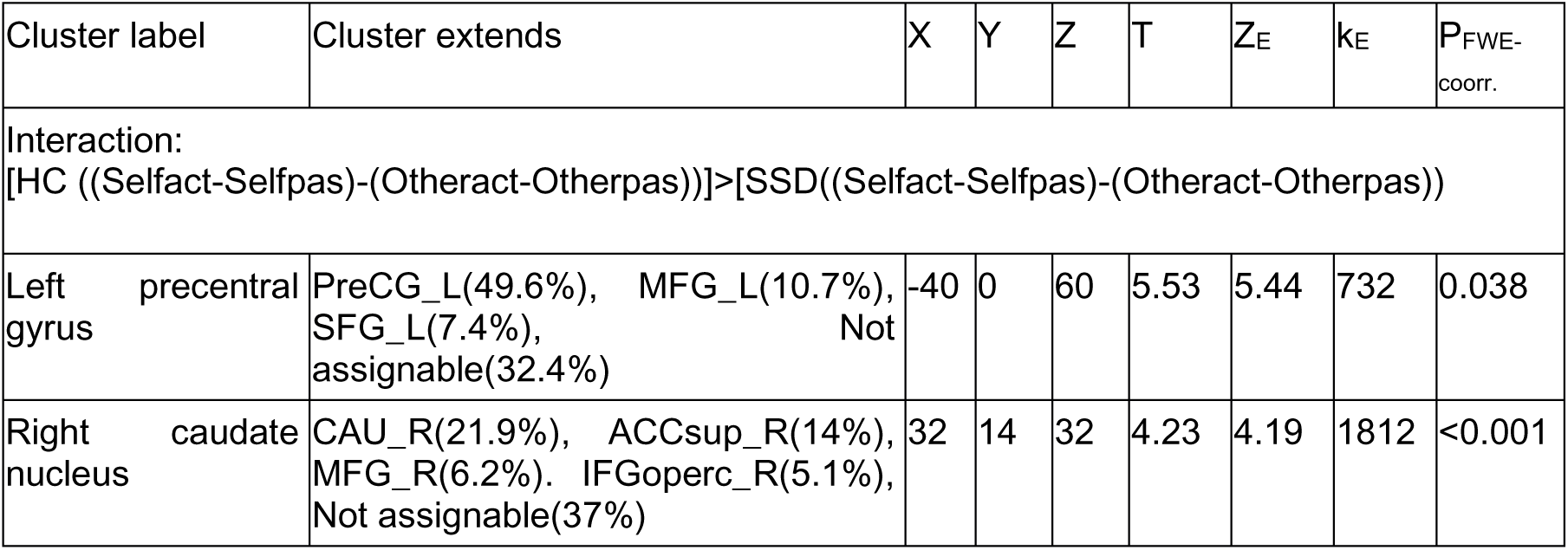
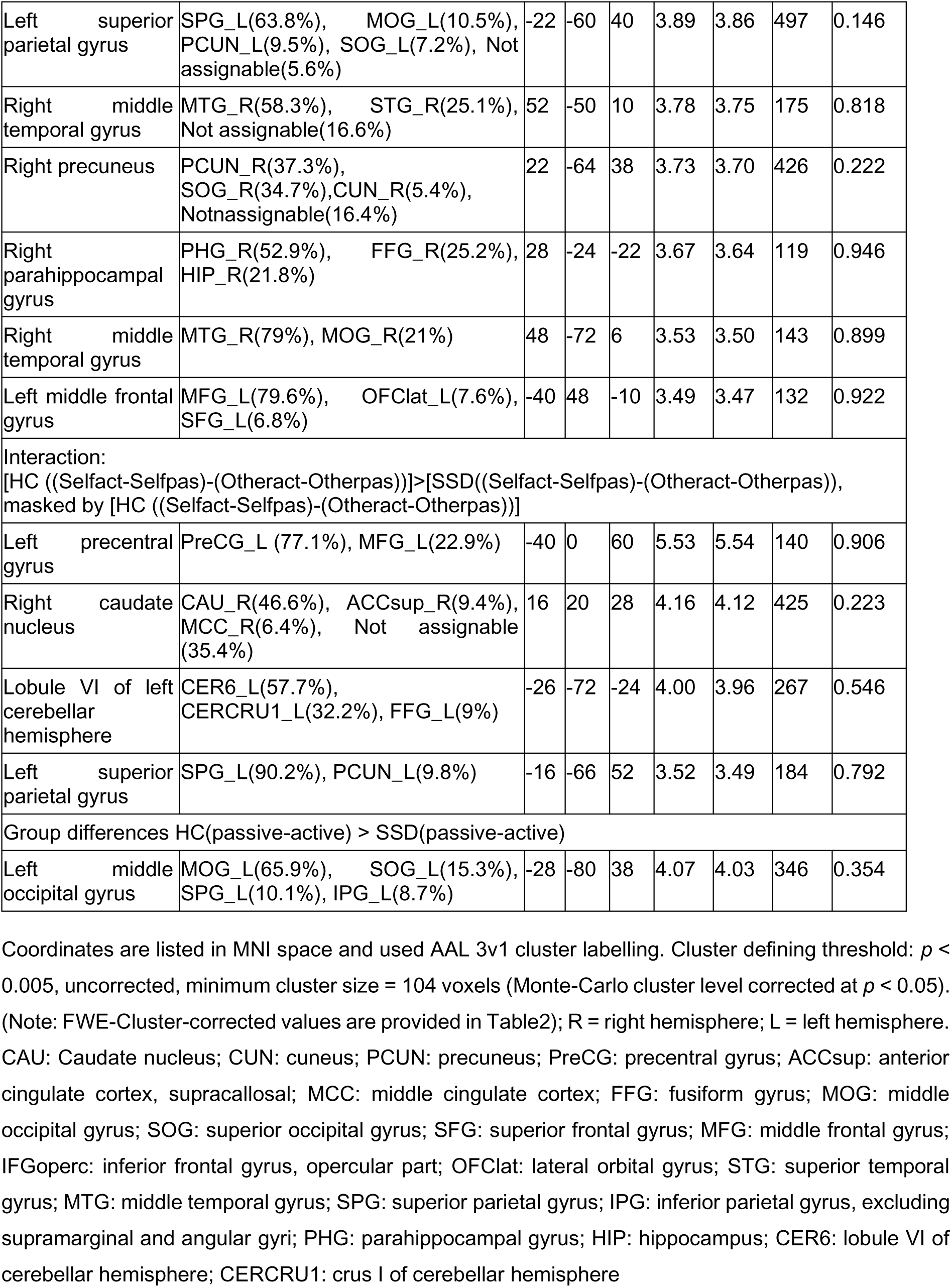
Results regarding DD: reflected activation duration in active compared to passive movement and vice versa.

**Figure 3.**
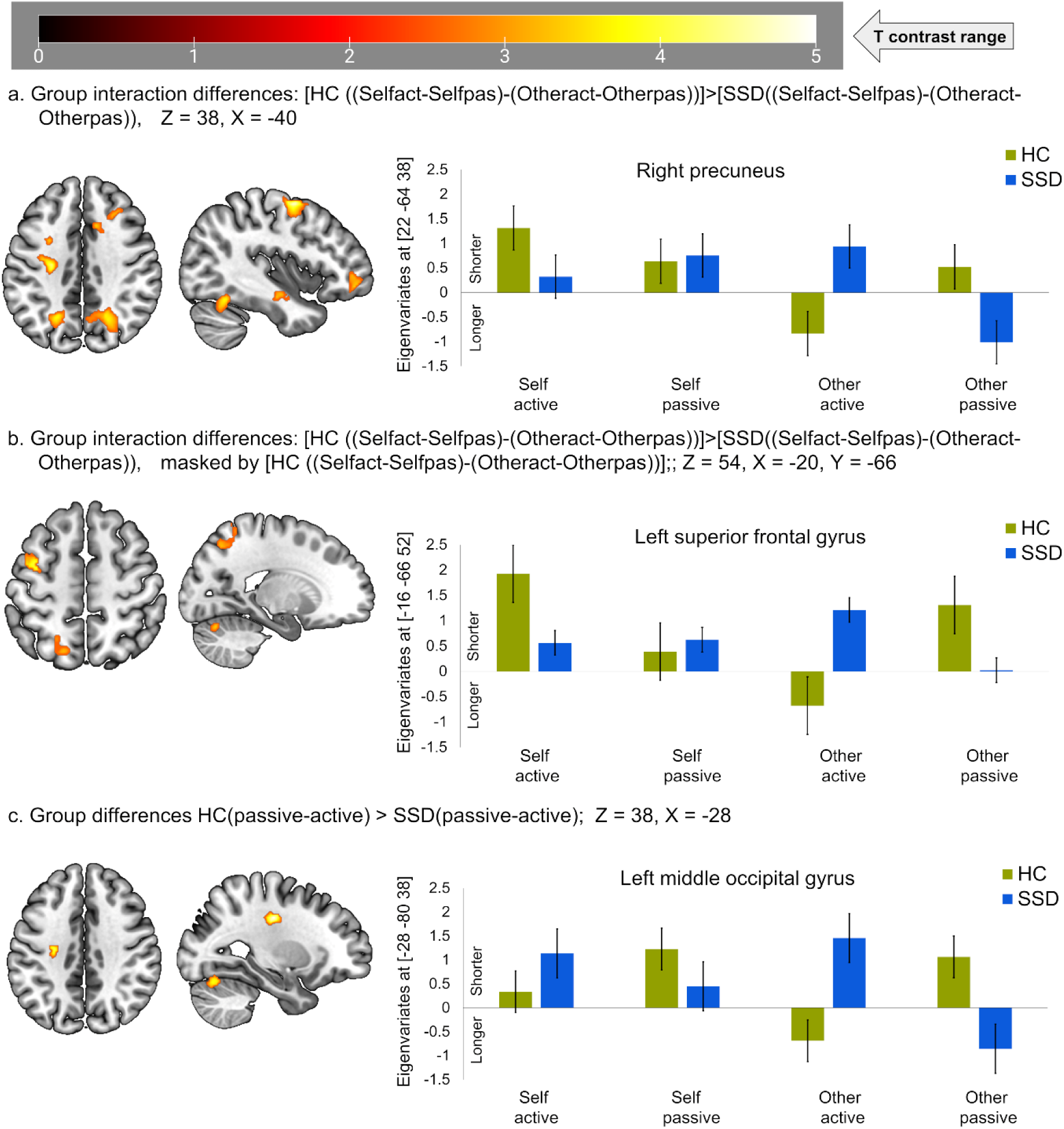
Duration of neural activation in active compared to passive and vice versa. (a) Group interaction: group (HC>SSD) X movement (active>passive) X feedback (own>other) hand shown at Z = 38, X = −40. b) Group interaction masked by HC X movement (active>passive) X feedback (own>other) shown at Z = 54, X = −20. (c) Group differences between passive and active movement regardless of feedback condition are shown at Z = 38, X = −28. Positive eigenvariates reflect shorter activation durations, while negative ones reflect longer activation durations in the bar graph. HC: healthy control, (n= 20); SSD: schizophrenia spectrum disorder, (n = 20).

**Figure 4.**
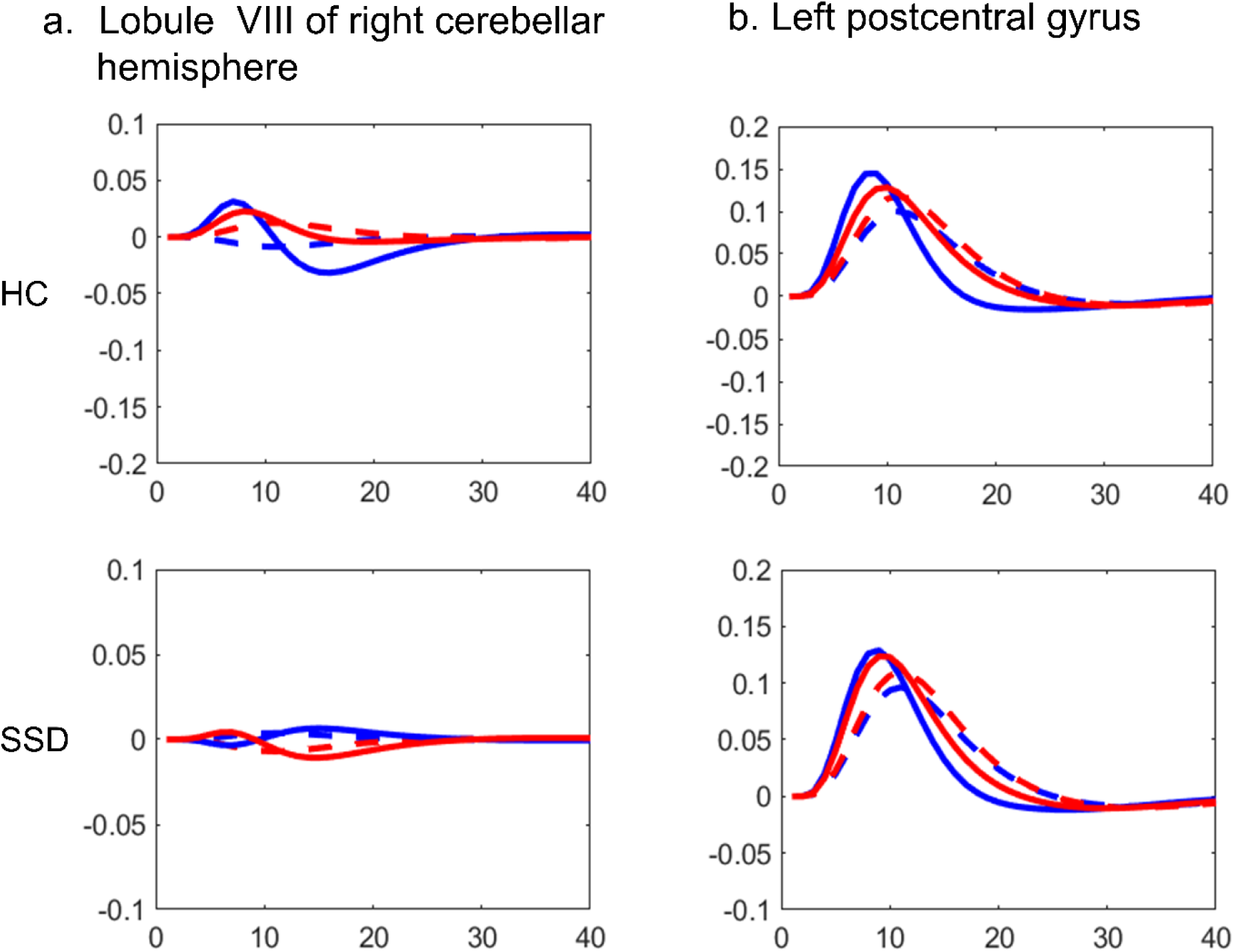
Latency functions (LF)^36,52^ from TD and from the canonical hemodynamic response function (HRF) in healthy control (upper row) and in schizophrenia spectrum disorder (SSD) in the lower row. (a.) LF at the lobule VIII of the right cerebellar hemisphere, and (b.) LF at the left post central gyrus. Active: canonical (dashed blue), HRF with temporal derivative (solid blue); Passive: canonical (dashed red), HRF with temporal derivative (solid red). HC: healthy control, (n= 20); SSD: schizophrenia spectrum disorder, (n = 20).

#### Duration of BOLD activation/suppression (passive>active)

The (passive > active) contrast did not reveal any regions with similar durations, nor any cluster for the interaction of group (HC>SSD), movement condition (passive > active), and video feedback (own hand > other hand). However, group differences (HC>SSD) with movement condition (passive > active) and video feedback of (own>other) hand revealed clusters comprising mainly the left middle occipital gyrus (Table 3, Figure 3c).

#### Exploratory correlation analyses

We explored the correlation between temporal dynamics of BOLD responses reflected in TD. Specifically, sub-cortical areas with earlier activation during active movement with own hand feedback (bilateral insula and putamen), and all positive and negative symptom scores were negatively related. A negative correlation indicates, that earlier processing (high or positive value for TD) is related to fewer symptoms, or the other way around: later processing (small or negative value for TD) is related to more symptoms. The timing of the left insula and putamen were negatively correlated to delusions of reference, delusions of being controlled, delusions, bizarre behaviour, residual positive symptoms, affective flattening or blunting, alogia, avolition/apathy, anhedonia/asociality, and total SANS score. The timing of the right insula and putamen was negatively correlated to delusions of reference, delusions of being controlled, delusions, residual positive symptoms, affective flattening or blunting, alogia, avolition/apathy, anhedonia/asociality, and total SANS score (Supplementary Table 1). However, partial correlation analyses controlling for total SANS score revealed no significant correlation (see Supplementary Table 2).

## Discussion

Here, we explored group differences and between group commonalities in the temporal dynamics of BOLD responses (considering the temporal derivative (TD) and dispersion derivative (DD)) for the processing of video feedback of active and passive hand movements. To our knowledge, this is the first fMRI study to identify aberrant temporal dynamics of BOLD responses during active and passive hand movements in patients with SSD. As hypothesised, we observed delayed neural processing for feedback of active movements in patients with SSD compared to HC in the right caudate nucleus, lobule VIII of the right cerebellar hemisphere, left superior temporal gyrus, left postcentral gyrus, left thalamus, and left putamen and insula. Delayed BOLD responses in bilateral insula were negatively correlated with positive and negative symptom severity. Additionally, unlike HC, patients with SSD did not show shorter processing in active compared to passive movement conditions with own hand feedback in the left precentral gyrus, right caudate nucleus, left superior parietal gyrus, and lobule VI of left cerebellar hemisphere. Furthermore, opposite to the HC, patients with SSD did not show shorter processing for passive movements in the left middle occipital gyrus. These changes in timing and duration may contribute to disrupted cortical-subcortical synchronization, sensory-motor integration, and impaired differentiation between active and passive movements. This impairment could potentially form the basis for impaired predictive mechanisms, disturbed sense of agency, and ego disturbances commonly observed in SSD. However, we could also state the main findings of commonalities. Patients and controls exhibited similar BOLD response timing in the clusters primarily comprising bilateral precentral gyrus, bilateral putamen, right fusiform gyrus, right middle temporal gyrus, right supramarginal gyrus, left supplementary motor area, left cuneus, and lobule VI of left cerebellar hemisphere (shown in Table 1, Figure 2c). These areas exhibited similar earlier BOLD responses in active versus passive hand movements, suggesting intact neural processing timing in these regions.

Previous research in HC has inferred that EC-based predictive mechanisms may lead to earlier neural processing of the visual feedback for active hand movement in contrast to passive ones.^36^ Advancing this background, we investigated how the temporal characteristics of BOLD responses for active and passive hand movements are deviating in patients with SSD compared to HC. Consistent with our hypothesis, we observed delayed neural processing for active movements feedback in patients with SSD compared to HC, including the right caudate nucleus, lobule VIII of the right cerebellar hemisphere, left superior temporal gyrus, left postcentral gyrus, left thalamus, and left putamen (Table 1, Figure 2d). However, interaction of [group (SSD > HC) X condition (active > passive) X feedback displayed (own hand > other hand)] further revealed that SSD showed relatively earlier activation (active>passive) in regions such as the left precentral gyrus, left supplementary motor area, and left postcentral gyrus (Table 1). The delayed activation in patients with SSD (Table 2, Figure 2d, 2e, and 2f) may stem from delayed and imprecise CD-mediated EC, leading to often reported delayed/disrupted and imprecise perception of sensory-motor feedback control (agency).^31,32,34,37,44^ This aligns with EEG studies suggesting that earlier activation-related RP may reflect intentional action awareness and motor preparation,^37,38^ manifested in earlier BOLD responses. These may indicate that the delayed BOLD response in SSD patients might be associated with a reduced advantage in predicting feedback from intended active versus passive hand movements.

We observed an earlier activation effect in the bilateral insula during active movements with own-hand feedback (Table 1, Figure 2f). The role of the insula is often reported as a core region of the self-network, linked to the feeling of causal or initiation of intended action, synchronisation of convergent internal-external sensory information, emotional context of action awareness, predictive processing of action-feedback, and intentional binding.^25,27,44,48,57–65^ Damage in the insula may manifest in disturbed self-action awareness in patients across psychiatric,^11,48,66^ neurological,^11,67^ and neurodegenerative disorders.^67–69^ Additionally, the neural activation timing in the bilateral insula was negatively correlated with positive and negative symptoms, excluding hallucinations, positive formal thought disorder, and attention, while the right insula further was not correlated to bizarre behaviour (Supplementary Table 1). Thus, impaired earlier activation timing of the bilateral insula extending to putamen in patients with SSD, especially during self-generated movement with own hand feedback, contributes to general symptom load rather than just self-other distinction or passivity symptoms.

Regarding the duration of BOLD responses, patients with SSD were mainly different from HC in the left precentral gyrus, right caudate nucleus, left superior parietal gyrus, right middle temporal gyrus, right precuneus, right parahippocampal gyrus, and left middle frontal gyrus as revealed by interaction analyses (Table 3, Figure 3a). However, when group interaction is masked by HC (Table 3, Figure 3a), patients with SSD demonstrated relatively longer processing for active movement with own hand feedback in the left precentral gyrus, right caudate nucleus, left superior parietal gyrus, and lobule VI of left cerebellar hemisphere. Although interaction effects are most distinct in active movement with own hand feedback, it is noticeable that durations of neural processing are altered in these brain areas in patients with SSD. Furthermore, regardless of the feedback conditions in the (passive > active) contrast, in contrast to HC, patients with SSD demonstrated longer processing for passive but shorter for active movements in the left middle occipital gyrus (Table 3, Figure 3c). These altered patterns of durations in these areas are mostly related to sensory-motor integration and monitoring active-passive movement authorship, especially with own hand feedback. ^8^

These findings with delayed timing (Table 1, Figure 2d, 2e, 2f) and altered durations of neural activation (Table 3, Figure 3a, 3b, 3c) suggest that patients with SSD may have impaired temporal BOLD response dynamics, leading to a disrupted timing and durations of cortical-subcortical synchronization. Temporally synchronised communication across brain regions across time is crucial during willed action awareness, thus their disruption likely contributes to the imprecise differentiation between active and passive movement authorship. In this regard, schizophrenia is characterised by movement abnormalities, excitation-inhibition imbalance, and delayed psychomotor activity may explain our altered temporal dynamics of BOLD responses in SSD patients.^3,46^ A review paper supports that psychomotor mediated neural activity patterns are abnormally balanced, and dynamic synchronisation across networks and brain regions is aberrant in schizophrenia.^70^ Further studies demonstrated that temporal variability of functional connectivity across time is disrupted in early visual areas, the thalamus, and the temporal cortex in patients with SSD.^71^ Another study reported abnormal dynamic changes in brain activity across time in schizophrenia.^72^ Thus, our findings may reflect a diminished dynamic ability to process awareness of intended hand movement, as seen in the impaired temporal dynamics of BOLD responses in patients with SSD.

Our findings largely align with the previous HC study^36^ regarding earlier activation during active compared to passive hand movements. Importantly, our findings further demonstrate aberrant patterns in SSD patients (Table 1, Figure 2d, 2f, 2e). However, some regions, particularly in SSD patients (Table 2) were less consistent with earlier activation during active than passive movement. Regarding durations of BOLD responses, shorter durations are more specific to active compared to passive movements, particularly with own hand feedback in HC, but this distinction was altered in SSD patients. These findings align with previous findings that patients with SSD exhibit impairments during preparation,^45^ which might explain the imprecise timing and durations of processing of intended movement related neural responses. Moreover, our timing results show that impairments may extend beyond movement preparation, revealing aberrant patterns of earlier-later BOLD response timing and altered durations during active and passive hand movements. These interpretations are supported by EEG studies of finger and brisk right fist closure movement in SSD patients, which demonstrated abnormal/altered action awareness, readiness potentials, and lateralised readiness potential amplitudes (reduced, delayed, mismatched in durations) typically emerge approximately 2s earlier than movement; along with deficits in post-movement event-related synchronisation.^73–76^ These abnormalities in temporal dynamics may reflect the sensorimotor integration abnormalities, potentially contributing to self-disorder in schizophrenia.^74^

Taken together, our findings emphasise the importance of considering the temporal dynamics of neural processing when examining sensory-motor dysfunctions in schizophrenia, which are undetectable by canonical analyses of BOLD amplitudes.^28^ The delayed and altered BOLD activation timing and distinct durations in SSD patients, along with the negative correlation of BOLD response timing to positive and negative symptom severity, may further confirm disruptions in the neural mechanisms underlying voluntary action and feedback monitoring. Understanding these abnormalities in temporal dynamics of BOLD responses provides key insights into the neural basis of sensory-motor dysfunction and self-disorder in patients with SSD.

### Limitations

Regarding generalisability, the relatively small sample size needs to be considered as a potential limitation, particularly in the correlation analyses. The application of statistical thresholds like family-wise error correction and correction for multiple comparisons has led to the disappearance of significance in most of these correlations. Patients were taking antipsychotics, we cannot rule out whether drug use might have some impact. In future experiments, large samples would be encouraged to map timing and durations in the whole brain. Furthermore, future studies could separate and investigate the timing for preparation and execution of hand movement, and how patients with SSD differ relative to HC. This could be an extension of previous findings of impaired preparatory neural activation patterns identified in patients with SSD.^45^ Although TD improves the detection of timing differences and DD captures changes in response duration, both can reduce sensitivity to amplitude-related effects.^53,54^ These trade-offs, inherent to basis set expansion, should be considered when interpreting group differences. Our sequential use of HRF+TD and HRF+TD+DD models aimed to balance these sensitivities, but some subtle effects may remain undetected depending on model choice. These complementary strengths and trade-offs should be carefully considered when interpreting results across models incorporating TD and DD.

## Conclusion

In this fMRI study, we investigated BOLD response timing using TD and durations using DD during the processing of active vs. passive hand movements with own or other hand video feedback. For the first time, we revealed commonalities and differences in BOLD response timing and durations between HC and patients with SSD. Our findings demonstrate aberrant temporal dynamics of BOLD responses in SSD patients, characterised by delayed and deviating BOLD response durations during active compared to passive hand movements. Our results suggest that these BOLD response characteristics may reflect deficits in the early predictive processing of schizophrenia. The distinct timing and duration pattern of the BOLD response across cortical and subcortical regions may indicate that temporal synchronization between areas is impaired, most likely due to imprecise or delayed corollary discharge and efference copy mechanisms. Such disruptions affect the temporal integration of internal and external signals, compromising the coordination needed for movement control and authorship. Notably, the reduced differentiation between active and passive movement-related BOLD responses in the bilateral insula-putamen and lobule VIII of the right cerebellar hemisphere, alongside the negative correlation between delayed BOLD responses in the insula-putamen and symptom severity, further highlight the link between these neural disruptions and disturbances in self-awareness and agency in schizophrenia. This study underscores the critical role of BOLD response timing and duration in understanding sensory-motor dysfunction in schizophrenia, offering new insights into the neural mechanisms underlying impaired action awareness, motor dysfunctions, and agency perception.

## Acknowledgements

We thank the Core Facility Brain Imaging Marburg and Lukas Uhlmann, Mareike Pazen, Anastasia Benedyk, Volker Besmens, Laila Noor, Lars Schwenzer, Jens Sommer, Olaf Steinsträter, and Dominik Vaughan for data collection as well as Jens Sommer for assistance with the implementation of the experimental setup.

## Supplementary material

See attached files

## Author contributions

Funding acquisition and project administration: Benjamin Straube. Conceptualization of the current study (manuscript): Harun A. Rashid, Benjamin Straube. Data analyses and results acquisition: Harun A. Rashid. Validation and supervision: Harun A. Rashid, Benjamin Straube. Writing – original manuscript: Harun A. Rashid. Further review and editing: Harun A. Rashid, Benjamin Straube, Tilo Kircher. All authors approved the manuscript.

