## Supplementary material for "Temporal characteristics of hemodynamic responses during active and passive hand movements in schizophrenia spectrum disorder": Supplenetary Table

### Supplementary material to the manuscript: Temporal characteristics of hemodynamic responses during active and passive hand movements in schizophrenia spectrum disorder

Supplementary Table 1. Correlation between the neural activation timing (cluster's eigenvariates) and symptom scores in SSD patients

| Cluster consisting mainly | symptom | Pearson correlation coeficient: r | p | Effect size (Fisher's z) | Spearman correlation coeficient: rho | p | Effect size (Fisher's z) |
|---|---|---|---|---|---|---|---|
| Left insula for active movement with own hand feedback | SAPS_I | -0.152 | 0.348 | -0.154 | -0.240 | 0.136 | -0.245 |
| | SAPS_14 | -0.517 | **<0.001** | -0.572 | -0.443 | 0.004 | -0.476 |
| | SAPS_15 | -0.475 | **0.002** | -0.516 | -0.443 | 0.004 | -0.475 |
| | SAPS_II | -0.500 | **0.001** | -0.549 | -0.491 | 0.001 | -0.537 |
| | SAPS_III | -0.317 | **0.046** | -0.329 | -0.296 | 0.064 | -0.305 |
| | SAPS_IV | -0.279 | 0.081 | -0.286 | -0.273 | 0.088 | -0.280 |
| | SAPS_res | -0.279 | **0.014** | -0.406 | -0.397 | 0.011 | -0.421 |
| | SANS_I | -0.419 | **0.007** | -0.447 | -0.335 | 0.035 | -0.348 |
| | SANS_II | -0.365 | **0.020** | -0.383 | -0.405 | 0.010 | -0.430 |
| | SANS_III | -0.536 | **<0.001** | -0.598 | -0.502 | <0.001 | -0.552 |
| | SANS_IV | -0.485 | **0.002** | -0.529 | -0.543 | <0.001 | -0.609 |
| | SANS_V | -0.085 | 0.602 | -0.085 | -0.088 | -0.591 | -0.088 |
| | SANS_total | -0.482 | **0.002** | -0.525 | -0.505 | <0.001 | -0.556 |
| Right insula for active movement | SAPS_I | -0.169 | 0.296 | -0.171 | -0.277 | 0.083 | -0.285 |
| | SAPS_14 | -0.485 | **0.002** | -0.525 | -0.430 | 0.006 | -0.459 |
| | SAPS_15 | -0.427 | **0.006** | -0.458 | -0.437 | 0.005 | -0.469 |
| | SAPS_II | -0.488 | **0.001** | -0.533 | -0.516 | <0.001 | -0.570 |

| | | | | | | | |
|---|---|---|---|---|---|---|---|
| with own | SAPS_III | -0.262 | 0.102 | -0.269 | -0.200 | 0.215 | -0.203 |
| hand | SAPS_IV | -0.241 | 0.133 | -0.246 | -0.298 | 0.061 | -0.308 |
| feedback | SAPS_res | -0.328 | **0.039** | -0.340 | -0.336 | 0.034 | -0.350 |
| | SANS_I | -0.388 | **0.013** | -0.409 | -0.332 | 0.037 | -0.345 |
| | SANS_II | -0.409 | **0.009** | -0.435 | -0.494 | 0.001 | -0.541 |
| | SANS_III | -0.478 | **0.002** | -0.520 | -0.441 | 0.004 | -0.474 |
| | SANS_IV | -0.474 | **0.002** | -0.516 | -0.535 | <0.001 | -0.597 |
| | SANS_V | 0.138 | 0.394 | 0.139 | 0.210 | 0.193 | 0.213 |
| | SANS_total | -0.468 | **0.002** | -0.507 | -0.466 | 0.002 | -0.504 |

Note: SAPS: scale for the assessment of positive symptoms, SAPS_I: hallucinations, SAPS_II: delusions, SAPS_14: delusions of reference, SAPS_15: delusions of being controlled, SAPS_III: bizarre behavior, SAPS_IV: positive formal thought disorder, SAPS_res: residual positive symptom (SAPS_III + SAPS_IV + SAPS_V), SANS: scale for the assessment of negative symptoms. SANS_I: affective flattening or blunting, SANS_II: alogia, SANS_III: avolition/apathy, SANS_IV: anhedonia/asociality, SANS_V: attention, SANS_total (SANS_I+SANS_II+SANS_III+SANS_IV+SANS_V). Bold values represent significant correlation ($p < 0.05$, uncorrected). A negative correlation indicates that earlier processing (high or positive value for temporal derivative (TD)) is related to less symptoms, or other was around later processing (small or negative value for TD) is related to more symptoms.

Supplementary Table 2 Partial correlation (partialling out SANS total score) between the neural activation timing (cluster's eigenvariates) and symptoms scores in SSD patients

| Cluster consisting mainly | symptom | Pearson correlation coeficient: r | p | Effect size (Fisher's z) | Spearman correlation coeficient: rho | p | Effect size (Fisher's z) |
|---|---|---|---|---|---|---|---|
| Left insula | SAPS_I | 0.201 | 0.220 | 0.204 | 0.071 | 0.669 | 0.071 |
| for active | SAPS_14 | -0.275 | 0.090 | -0.282 | -0.170 | 0.301 | -0.172 |
| movement | SAPS_15 | -0.212 | 0.196 | -0.215 | -0.182 | 0.268 | -0.184 |
| with own | SAPS_II | -0.151 | 0.359 | -0.152 | -0.051 | 0.760 | -0.051 |
| | SAPS_III | -0.253 | 0.121 | -0.258 | -0.190 | 0.248 | -0.192 |

| | | | | | | | |
|---|---|---|---|---|---|---|---|
| hand feedback | SAPS_IV | -0.110 | 0.505 | -0.111 | -0.088 | 0.594 | -0.088 |
| | SAPS_res | -0.219 | 0.180 | -0.223 | -0.239 | 0.143 | -0.244 |
| Right insula for active movement with own hand feedback | SAPS_I | 0.163 | 0.322 | 0.164 | -0.014 | 0.932 | -0.014 |
| | SAPS_14 | -0.231 | 0.157 | -0.235 | -0.185 | 0.260 | -0.187 |
| | SAPS_15 | -0.149 | 0.367 | -0.150 | -0.206 | 0.208 | -0.209 |
| | SAPS_II | -0.156 | 0.341 | -0.158 | -0.261 | 0.109 | -0.267 |
| | SAPS_III | -0.191 | 0.245 | -0.193 | -0.086 | 0.605 | -0.086 |
| | SAPS_IV | -0.070 | 0.672 | -0.070 | -0.137 | 0.406 | -0.138 |
| | SAPS_total | -0.154 | 0.350 | -0.155 | -0.178 | 0.280 | -0.179 |

Note: SAPS: scale for the assessment of positive symptoms, SAPS_I: hallucinations, SAPS_II: delusions, SAPS_14: delusions of reference, SAPS_15: delusions of being controlled, SAPS_III: bizarre behavior, SAPS_IV: positive formal thought disorder, SAPS_res: residual positive symptom (SAPS_III + SAPS_IV + SAPS_V), SANS: scale for the assessment of negative symptoms. SANS_I: affective flattening or blunting, SANS_II: alogia, SANS_III: avolition/apathy, SANS_IV: anhedonia/asociality, SANS_V: attention, SANS_total (SANS_I+SANS_II+SANS_III+SANS_IV+SANS_V). Bold values represent significant correlation ($p < 0.05$, uncorrected).

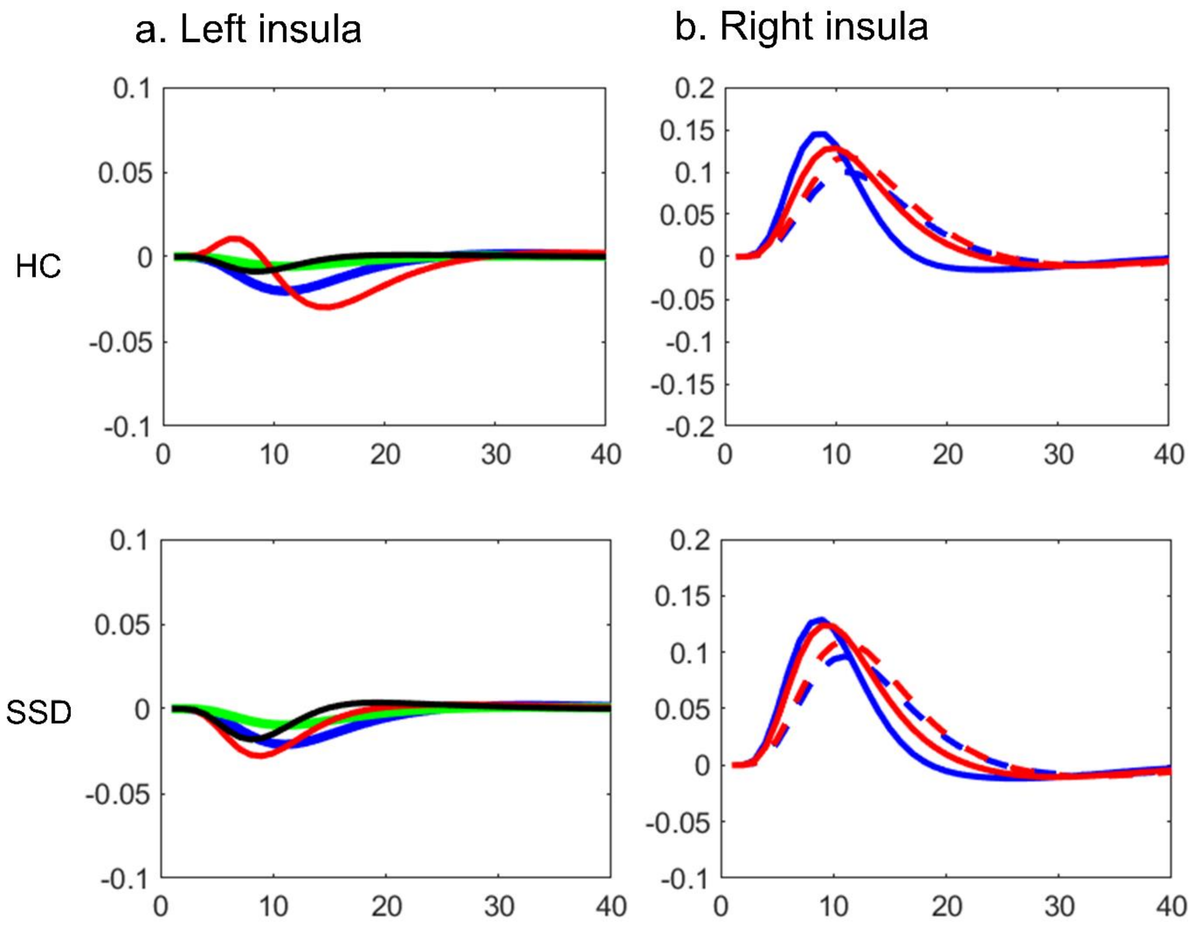


Supplementary Figure 1 Timing in insula during active movement with own hand feedback. Latency functions (LF) from DD and from the canonical hemodynamic response function (HRF) in healthy control (upper row) and in schizophrenia spectrum disorder (SSD) in the lower row. (a.) LF at the left insula, and (b.) LF at the right insula. Active: canonical (dashed blue), temporal derivative (solid blue); Passive: canonical (dashed red), temporal derivative (solid red).
